# Deriving Cardiomyocytes from Human Amniocytes

**DOI:** 10.1101/475624

**Authors:** Colin T. Maguire, Ryan Sunderland, Bradley Demarest, Bushra Gorsi, Josh Jackson, Angelica Lopez-Izquierdo, Martin Tristani-Firouzi, H. Joseph Yost, Maureen L. Condic

**Affiliations:** University of Utah Molecular Medicine Program (U2M2); Department of Neurobiology and Anatomy; Nora Eccles Harrison Cardiovascular Research & Training Institute; Department of Pediatrics, University of Utah, School of Medicine, 30 N. 1900 E. Salt Lake City, UT 84132, USA

**Author notes:** Correspondence: Colin T. Maguire, PhD, Cellular Translational Research Core, Utah CCTS, 26 N Medical Drive, Wintrobe, Rm. 203, Salt Lake City, Utah 84112.

**Keywords:** Amniotic fluid, Amniocyte, Induced-pluripotent stem cells, Cardiomyocyte differentiation, Methylome, Transcriptome.

## Abstract

Many forms of congenital heart disease (CHD) have high morbidity-mortality rates and require challenging surgeries. Human amniocytes have important stem cell characteristics and could potentially provide patient-specific tissue for repairs of some types of CHDs. We report that amniocytes express features of poised cardiomyocytes. However, a variety of direct reprogramming approaches failed to convert their fetal and transcriptionally repressed state into *bona fide* cardiomyocytes. Induced-pluripotent stem cell (iPSC) reprogramming removes repression and converts amniocytes to a baseline pluripotent state. Based on molecular and electrophysiological signatures, iPSC reprogrammed amniocytes can be induced to differentiate into functionally immature, predominantly ventricular cardiomyocytes and a heterogeneous mixture of vascular and unspecified epithelial cells. Developmental time course analyses and pattern clustering of amniocyte-derived cardiomyocytes identifies numerous temporal co-regulators of cardiac induction and maturation as well as distinct sarcomeric and ion channel gene signatures. Normal fetal cardiomyocytes are derived by overcoming complex forms of transcriptional repression that suppress direct transdifferentiation of human amniocytes. These results suggest the possibility of using amniocytes as a source of patient-specific ventricular cardiomyocytes for cell therapies.

**SUMMARY STATEMENT:** Amniocytes are a possible source of patient-specific cardiomyocytes for newborns with congenital heart disease. Genome-wide DNA methylation patterns and transcriptional repressors preclude direct differentiation, but pluripotent reprogramming provides cardiomyocytes for dissecting genetic pathways contributing to this disease.

## INTRODUCTION

Congenital Heart Disease (CHD) is a serious and common condition that claims more children’s lives than any other human birth defect (Benjamin et al., 2017; Hoffman and Kaplan, 2002). Some forms of CHD require surgery in the first weeks or months after birth. Since earlier postnatal interventions reduce myocardial damage and reduce mortality (Gardiner, 2009), earlier fetal interventions are being sought (Freud and Tworetzky, 2016; Marantz and Grinenco, 2015; McLaughlin et al., 2016; Tulzer and Arzt, 2013). Stem-cell based therapies and engineered cell sheets may offer a new line of treatment, in conjunction with standard surgical approaches. From a bioethical, technical, and economic standpoint, reprograming cells with chemical enhancement (Abad et al., 2017; Mohamed et al., 2017) represents the best option for most clinical researchers. Currently, at least eight potential sources of cardiac specific stem cells are being considered for transplantation (Bernstein and Srivastava, 2012), including amniotic-fluid derived cells (Kunisaki, 2018).

Amniocytes are fetal cells, developmentally linked to fetal maturation (Dobreva et al., 2010). They are easily obtainable from standard diagnostic amniocentesis as early as 15 weeks of gestation, with minimal risk to the mother or fetus (Corrado et al., 2012; Lenis-Cordoba et al., 2013). This means amniocytes represent one of the earliest autologous cell-types that could be obtained, manipulated *ex-utero* and surgically returned to the same patient, without facing immune rejection. Amniocytes are highly proliferative when grown *in vitro*, but they do not produce tumors *in vivo*, and are not immortal (De Coppi et al., 2007). Since amniocytes reside in a developmentally immature state, they retain a certain level of innate pluripotency. In fact, amniocytes are more efficiently reprogrammed back to a pluripotent state than postnatal- or adult-derived cell-types (Easley et al., 2012; Galende et al., 2009; Li et al., 2009). Genome-wide studies in our lab suggest amniocytes express a distinct multilineage phenotype with pluripotent features (Maguire et al., 2013), making them an intriguing candidate for reprogramming and transplantation studies. Recent direct reprogramming approaches using defined combinations of reprogramming factors, microRNAs, small-molecules and cardiac-inductive signals have converted mouse fibroblasts (Efe et al., 2011; Ieda et al., 2010; Jayawardena et al., 2012; Qian et al., 2012; Song et al., 2012; Wang et al., 2014) and human fibroblasts (Cao et al., 2016; Fu et al., 2013; Nam et al., 2014; Wada et al., 2013) into functional cardiomyocytes (Zhang et al., 2016).

Given that amniocytes possess some pluripotent features, we tested the hypothesis that amniocytes would be readily amenable to direct differentiation, using a battery of small molecules, cardiac-inductive signals, and minicircle reprogramming approaches. Despite having innate early cardiac characteristics, amniocytes were unexpectedly resistant to direct cardiomyocyte conversion, even using protocols previously successful in fibroblasts. While resistant to direct differentiation protocols, amniocytes were amenable to reprogramming back to a pluripotent state by delivery of standard reprogramming genetic factors, and then efficiently differentiated into a fetal cardiovascular fate. Analysis of the temporal expression patterns of amniocytes converted to cardiomyocytes allowed the identification of potential coregulators of cardiovascular differentiation.

## RESULTS

### Amniocytes are Poised Cardiomyocytes

The human cardiac transcriptome has been published from individuals at different ages (Asp et al., 2012; Nanni et al., 2006; Sanoudou et al., 2005) but cardiac transcription has not been extensively studied across developmental time. In order to set a baseline transcriptional signature, we combined multiple publicly available RNA-seq datasets from the human heart at three different stages of life (fetal, children, and adult). Groups of genes known to play a role in early-, mid-, and late-cardiac development express a transcriptional signature that is remarkably similar among fetal, children, and adult hearts (Figure 1A, panel of 57 cardiac markers). Strikingly, this analysis indicates that once the embryonic heart forms (8 weeks of gestation), important cardiovascular markers and pathways are maintained throughout maturation of the fetus, during childhood and during adulthood.

**Figure 1.**
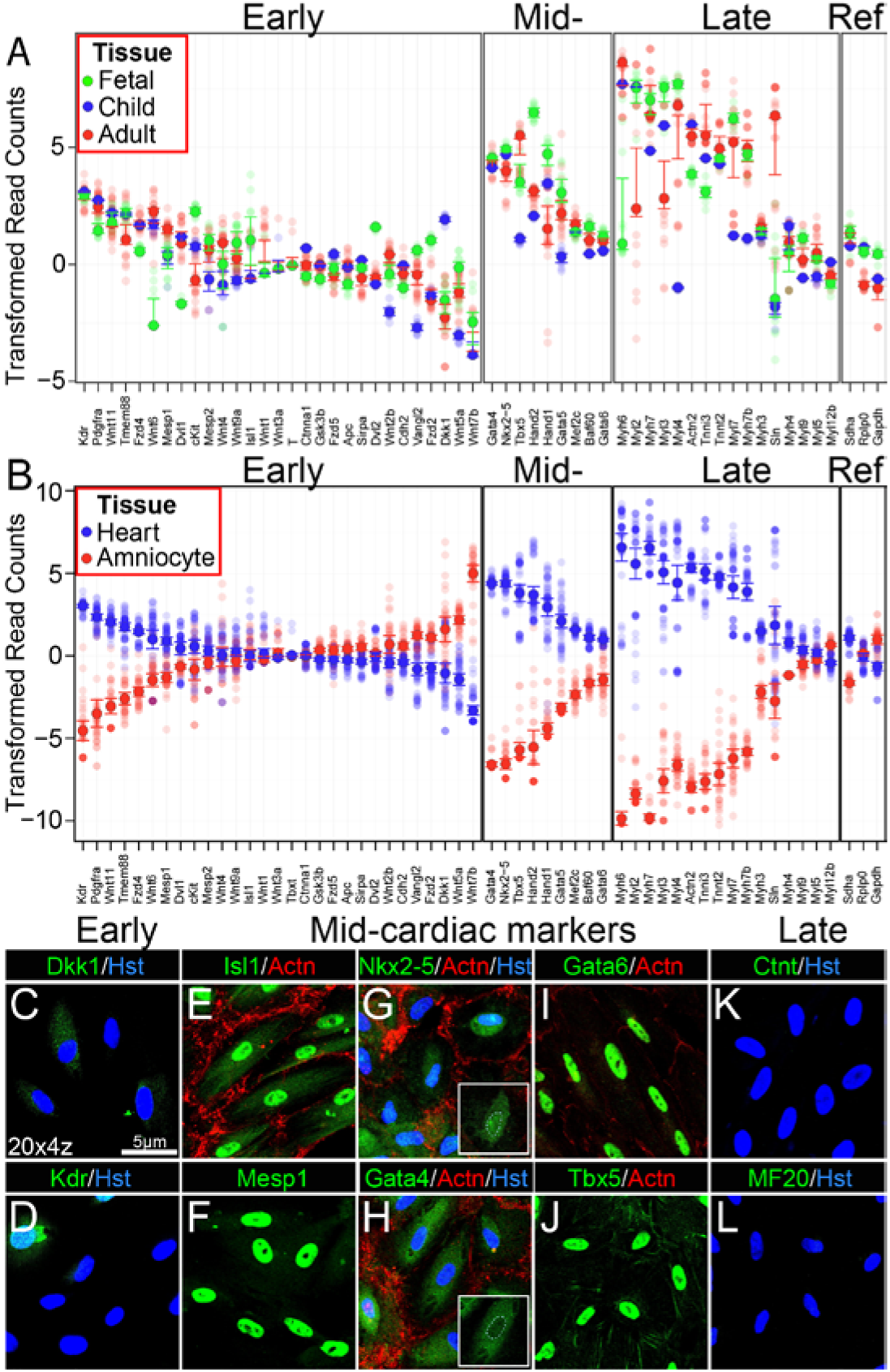
Amniocytes are Poised Cardiomyocytes. (A): RNA-seq scatterplot comparing fetal, child, and adult gene expression in human hearts for selected early-, mid-, and late-cardiac markers. Green dots = fetal hearts, blue dots = children hearts, and red dots = adult hearts (N= 48 RNAseq samples). Solid circles are the median value (bars ± standard deviation) for each group and the transparent circles are expression levels for individual samples. (B): RNA-seq scatterplot comparing heart and amniocyte gene expression for selected early-, mid-, and late-cardiac markers. Blue dots = human heart and red dots = human amniocytes. Solid circles are the median value (bars ± standard deviation) for each group and the transparent circles are expression levels for individual samples. (C-L): Immunofluorescence analysis of (C and D) early-, (E, F, G, H, I, and J) mid-, and (K and L) late-cardiac markers. We examined 21 different amniocyte patient samples (Dkk1 N = 3, Kdr N = 3, Isl1 N = 8, Mesp1 N = 3, Nkx2-5 N = 5, Gata4 N = 10, Gata6 N = 5, Tbx5 N = 7, Ctnt N = 4, MF20 N = 1). (G and H) insets without nuclear counterstain showing no nuclear staining of two key core cardiac transcription factors. Hst = Hoechst dye nuclear counterstain in b

The developmental status of amniocytes remains unclear. Ongoing genome-wide studies, qPCR confirmation, and immunofluorescence studies in our lab indicate amniocytes display an unusual cellular phenotype, with multiple germ-layer specific regulators being co-expressed (Maguire et al., 2013). Here, we focus on the mesodermal lineage and examine whether amniocytes express a genes indicative of a cardiac signature. Confocal images of immunostained amniocytes indicate expression of early cardiac markers at low levels (Figure 1C and 1D), as well as some mid-phase cardiac markers (Figure 1E,1F, 1G, and 1H). Strikingly, cardiac transcription factors Nkx2-5 and Gata4 are expressed but are localised to the cytoplasm, not to the nucleus (Figure 1G and 1H). Markers of full cardiomyocyte differentiation, for example sarcomeric proteins, are undetectable (Figure 1K and 1L). This data indicates amniocytes have some characteristics of cardiac progenitors or early cardiomyoctes, but the later stage cardiomyocyte transcriptome is repressed.

Since amniocytes express some early-and mid-phase cardiomyocyte markers, we examined how closely related amniocytes are to heart tissues on a genome-wide level. Genome-wide scatterplots reveal amniocytes and heart tissue have a weak relationship with a correlation coefficient of 0.52 (Sup Figure 1). Using a panel of early-, mid-, and late-cardiac markers, expression of most of the mid- and late-cardiac markers was markedly reduced in amniocytes compared to heart tissue (Figure 1B). Nonetheless, amniocytes not only show some similarities to heart in genome-wide expression, but they express important developmental cardiac markers, suggesting amniocytes exhibit features of poised cardiomyocytes. This suggests that amniocytes have progressed part-way along cardiovascular developmental pathways but are either lacking activity of key transcriptional factors (as confirmed for Nkx2.5 and Gata4, Figure 1G and 1H) and/or express inhibitors of these pathways.

### Amniocytes are resistant to direct differentiation into cardiovascular lineages

Given that amniocytes possess some pluripotent features, we tested the hypothesis that amniocytes would be readily amenable to direct differentiation. To determine whether the amniocyte transcriptome could be pushed toward a cardiomyocyte fate, we first manipulated cardiogenic pathways using different combinations of cardiomyocyte-inducing small molecules, recombinant signaling factors, and chromatin modifying agents as well as commercial cardiomyocyte differentiation kits and published cardiomyocyte differentiation protocols (Burridge et al., 2014; Efe et al., 2011; Lian et al., 2012; Yang et al., 2008; Zhang et al., 2012) and monitored cardiomyocyte differentiation using a panel of 19 cardiovascular markers (Sup Figure 2). In order to better understand if amniocytes can be efficiently differentiated, we used non-integrating minicircles to knock down the most highly expressed embryonic repressors to determine whether any of them (singly or combinations) contribute to repression of amniocyte differentiation toward a cardiac lineage. As an example, transfection of Gata4-containing minicircles into amniocytes resulted in nuclear localised Gata4 protein and consistently increased gene expression over 500 fold, as measured by both qPCR and RNA-seq (Sup Figure 3). Yet surprisingly, even overexpression of key cardiogenic factors together with simultaneous siRNAs knockdown of specific cardiac inhibitors [21] did not result in full differentiation of amniocytes into cardiomyocytes (Sup Figure 4).

**Figure 2.**
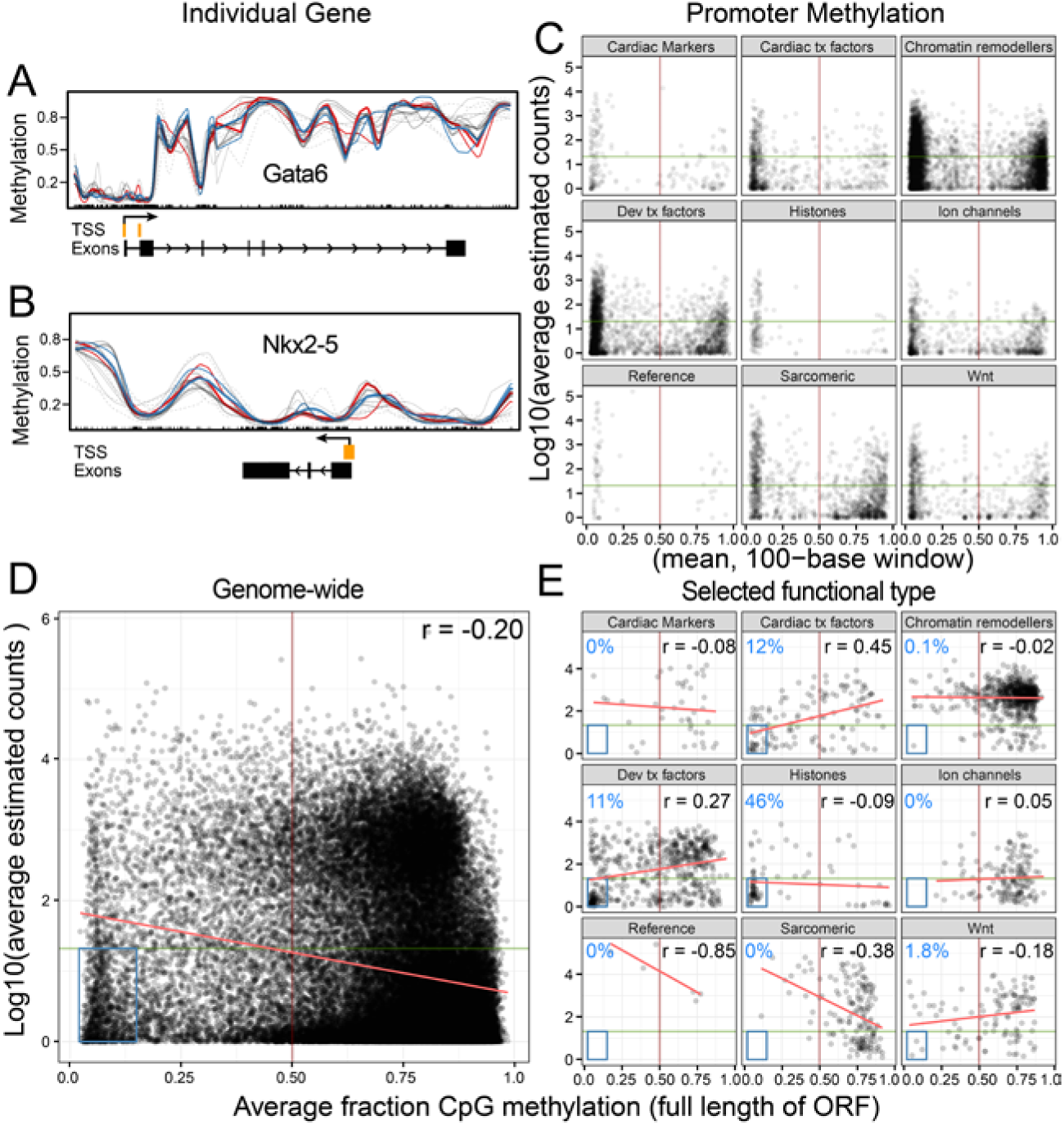
Amniocyte DNA methylation and gene expression is nonconforming. (A): BSsmooth plot derived from GWBS for Gata6. Based on DNA methylation, qPCR, RNA-seq, and immunostaining data, Gata6 is an example of an expressed cardiac-specific gene that has a hypomethylated TSS and a hypermethylated ORF (ENSG00000141448, + strand, chr18: 19,749,404 − 19,782,491 (width = 33,088, extended = 5,000). Blue = Control, DMSO-treated, high sequencing coverage. Red = 5-Aza. agents, high sequencing coverage. Grey = low coverage. (B): In contrast, BSsmooth plot derived from GWBS for Nkx2-5. Based on DNA methylation, qPCR, RNA-seq, and immunostaining data, Nkx2-5 is an example of a cardiac-specific gene not expressed that has a hypomethylated TSS and a hypomethylated ORF (ENSG00000183072, − strand, chr5: 172,659,112 − 172,662,360 (width = 3,249, extended = 5,000). (C): Genome-wide scatterplot of amniocyte DNA methylation and gene expression. Gene expression derived from 37 RNA-seq samples transformed into Log10 (average estimated counts). For each selected gene’s open reading frame, GWBS data was used to calculate the average fraction of CpG methylation. (D): The nine panels in (D) have been organised by functional gene type or pathway. Note the percentage of TTS in the blue box and regression slope vary depending on gene functional type or pathway. The vertical brown line denotes the boundary separating high and low methylation (0.50 cutoff), while the horizontal green line denotes the boundary separating high and low gene expression (20 read cutoff). The red line is the linear regression slope. The blue number located on the top left of each panel is the percentage of TSSs that fall inside the blue box. The black number located on the top right of each panel is the Pearson’s r correlation value.

**Figure 3.**
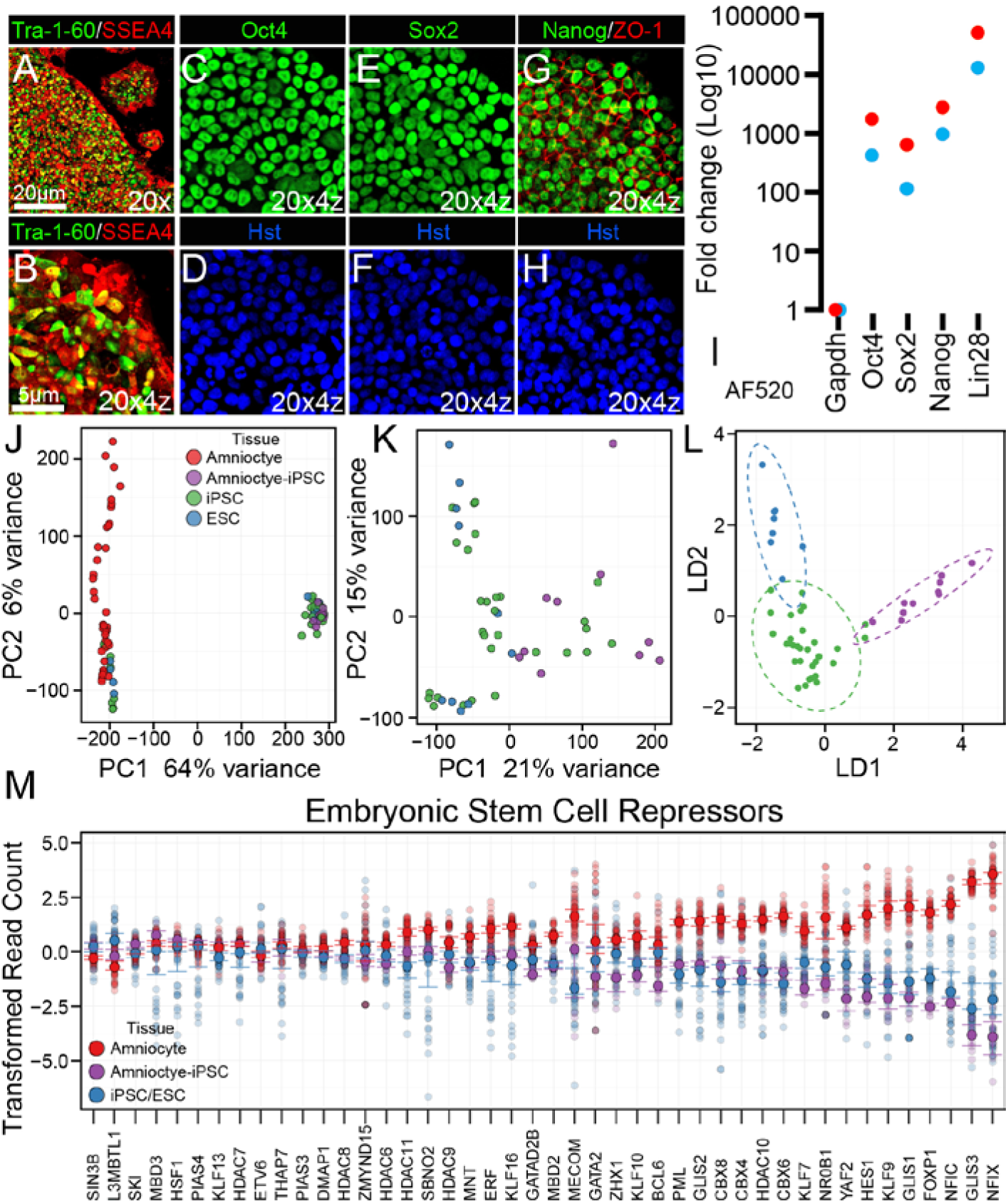
Amniocyte-iPSCs overcome repression and convert into a typical pluripotent state. (A-H): Confocal immunostaining of newly formed amniocyte-iPSCs (N = 6 patient A-iPSC lines) (A and B): co-staining pluripotent surface markers. (C and E): pluripotent nuclear markers Oct4 and Sox2. (G): Nuclear pluripotent marker Nanog and epithelial marker ZO-1. (D, F, and H) Nuclear counterstain. (I): qPCR analysis of Gapdh (reference) and pluripotent makers showing dramatic increases in expression for Oct4, Sox2, Nanog, and Lin28. Amniocyte-iPSCs were grown on victronectin-(red dots) and matrigel-(blue dots) coated plates. (J): Principal component analysis of unmanipulated amniocytes (red), A-iPSCs (violet) and previously published iPSCs (aqua) and human embryonic stem cell lines (green). In panel J, A-iPSCs (N = 10), iPSCs (N = 49) and ESCs (N = 19) cluster closely together, making it difficult to examine any similarities or differences. (K) In order to better analyze the overlapping cluster of stem cell lines, unmanipulated amniocytes were excluded. (L): Linear discriminant analysis of A-iPSCs and previously published iPSCs and stem cell lines. (M): 44 embryonic stem cell repressors expressed at high levels in unmanipulated amniocytes (red, N= 37). A-iPSCs (purple, N= 10) reduce expression of these 44 repressors to match expression levels of other pluripotent cell lines (blue, N= 68).

**Figure 4.**
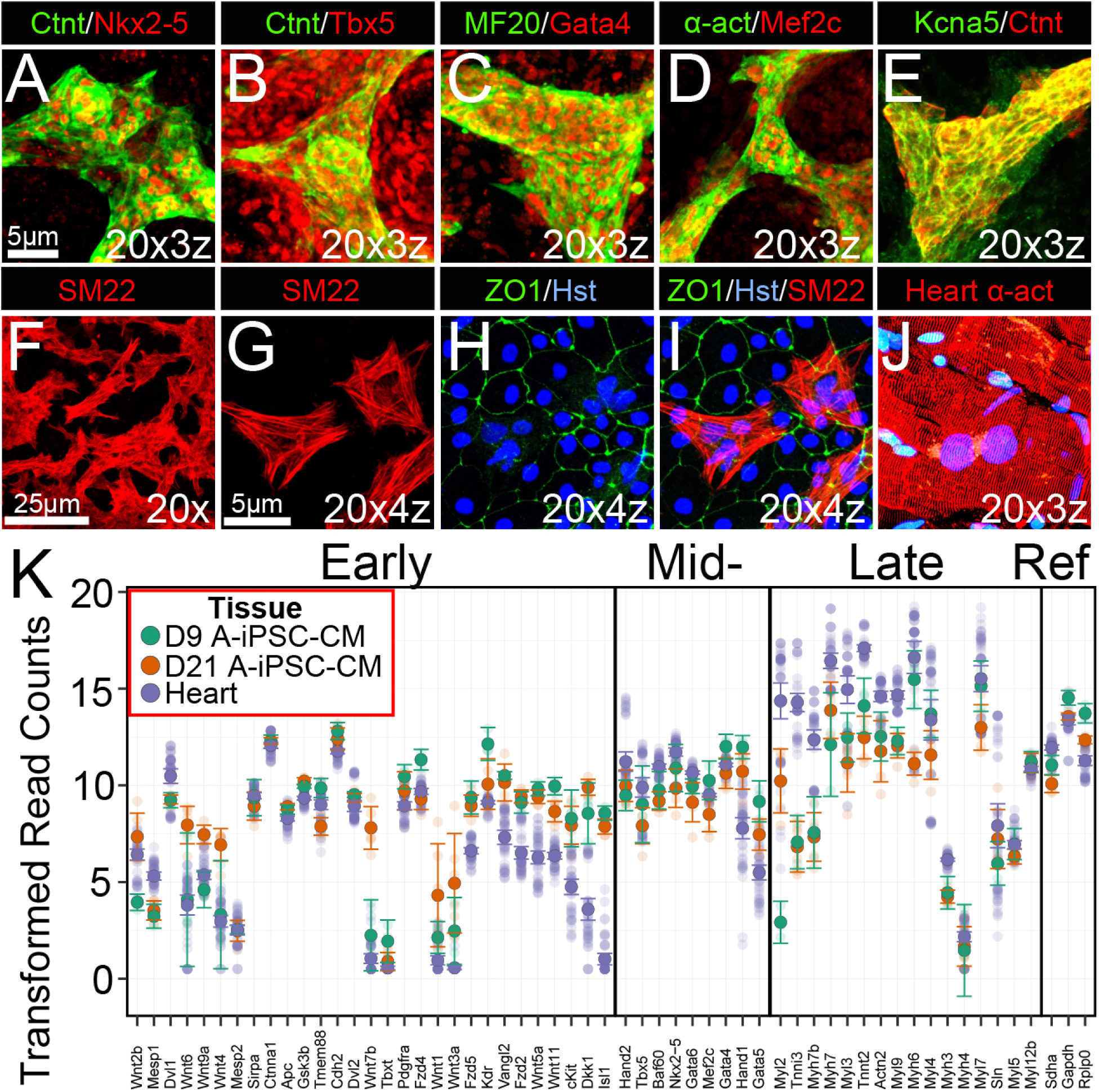
Amniocyte-iPSC-cardiomyocytes match a human phenotype. (A-E): Cultured A-iPSC-CMs immunostained with cardiac markers. (F-I): Cultured A-iPSC-cardiovascular cells immunostained with smooth muscle marker SM22 and epithelial marker ZO-1, nuclear counterstained (blue) with Hoechst dye. (J): Adult human heart tissue immunostained with alpha-actinin and nuclei counterstained (blue) with Hoechst dye. (K): RNA-seq scatterplot comparing human heart (purple, N= 48) to induction A-iPSC-CMs (Day9, green, N = 4) and maturation A-iPSC-CMs (Day21, blue, N = 6). Selected genes were grouped by early-, mid-, and late-cardiac markers. Solid circles are the median value (bars ± standard deviation) for each group and the transparent circles are expression levels for individual samples.

To assess whether Gata4 overexpression altered expression levels of its target genes in amniocytes, we curated 122 putative Gata4 target genes from literature (Sup Table 2). Surprisingly, only four Gata4 target genes increased significantly in Gata4 overexpressing amniocytes, and the expression levels for most known Gata4 target genes remained unchanged (Sup Figure 5). While Oas2 (2’-5’-oligoadeny-late synthetase 2), along with other Oas family members increased their levels, these genes are unlikely to play a role in cardiovascular differentiation. Thus, while Gata4 is a pioneer transcription factor for cardiogenesis (Cirillo et al., 2002; Zaret and Carroll, 2011), the inability to activate cardiovascular gene programs in Gata4 overexpressing amniocytes suggests that target transcription binding sites are inaccessible.

**Figure 5.**
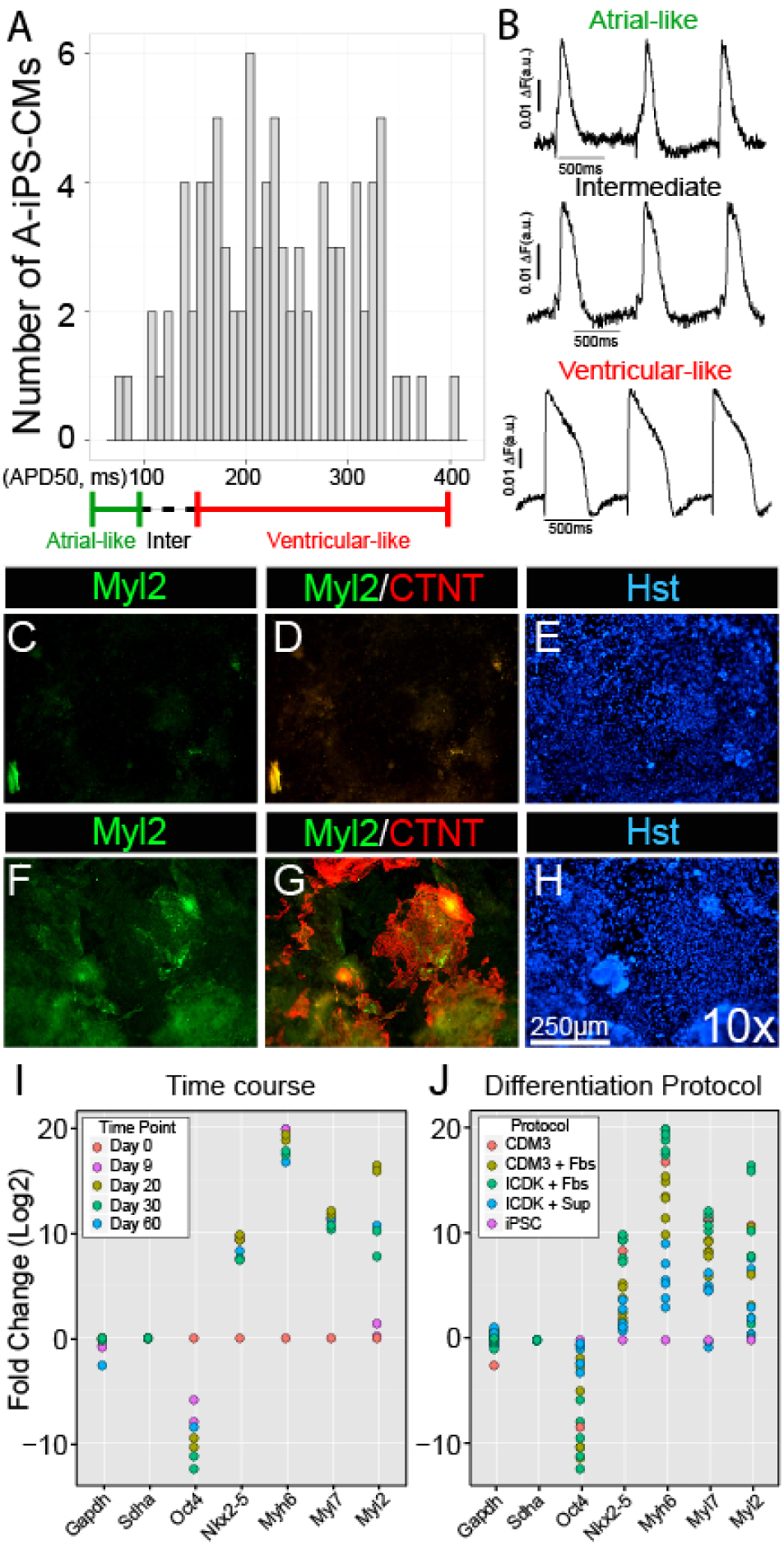
Most A-iPSC-CMs are ventricular-like cardiomyocytes. (A): A bar plot of electrophysiological recordings from individual contracting A-iPSC-CMs (N = patient sample). Atrial-like cells have APD50s up to 100ms, ventricular-like cells have APD50s ranging from 150ms-400ms, and cardiomyoctes falling between these two groups are designated as an intermediate phenotype. (B) Action potential recordings from atrial-like, intermediate, and ventricular-like A-iPSC-CMs. (F-H) Day 20 A-iPSC-CM contracting sheets immunostained with the cardiomyocyte marker CTNT (red) and ventricular marker MLC2V (Myl2) (green), nuclear counterstained (blue) with Hoechst dye. (I) qPCR time course of CDM3 cardiomyocyte differentiation protocol at Days 0, 9, 20, 30 and 60. qPCR analysis of Gapdh and Sdha are reference genes, Oct4 is a pluripotent marker, Nkx2-5 is a cardiac transcription factor, Myh6 is a cardiac sarcomeric gene, Myl7 is an atrial marker, Myl2 is a ventricular marker. (J) qPCR analysis of contracting A-iPSC-CMs at Day 20 using two different cardiomyocyte differentiation protocols with or without additional culture supplements or FBS.

In addition to minicircle-Gata4, forced expression of a mesoderm activator (minicircle-T/Brachyury-GFP) and cardiogenic transcription factors (minicircle-Nkx2-5-GFP and minicircle-Tbx5-GFP) did not significantly alter expression of the transcriptional network of cardiogenic genes. In unmanipulated amniocytes, qPCR analysis could not detect any cardiogenic makers above baseline. At 24 hours post-transfection, minicircle-overexpressed transcription factors were significantly elevated, but downstream key core transcription factors and other cardiogenic genes on our cardiac panel remained essentially unchanged. These results suggest that using minicircle technology to force expression of cardiac transcription factors (individually or in combination) may not be a powerful enough overexpression system to efficiently reprogram human amniocytes into cardiogenic pathways.

Since amniocytes have innate pluripotent characteristics, we also tested whether the small molecule valproic acid or minicircle-LGNSO (Lin28, GFP, Nanog, Sox2, Oct4) could reprogram amniocytes to a partial or full pluripotent state. Using valproic acid to chemically-induce amniocytes into pluripotency (Hawkins et al., 2017; Moschidou et al., 2012), key pluripotent markers decreased, suggesting this option was insufficient (Sup Figure 6).

**Figure 6.**
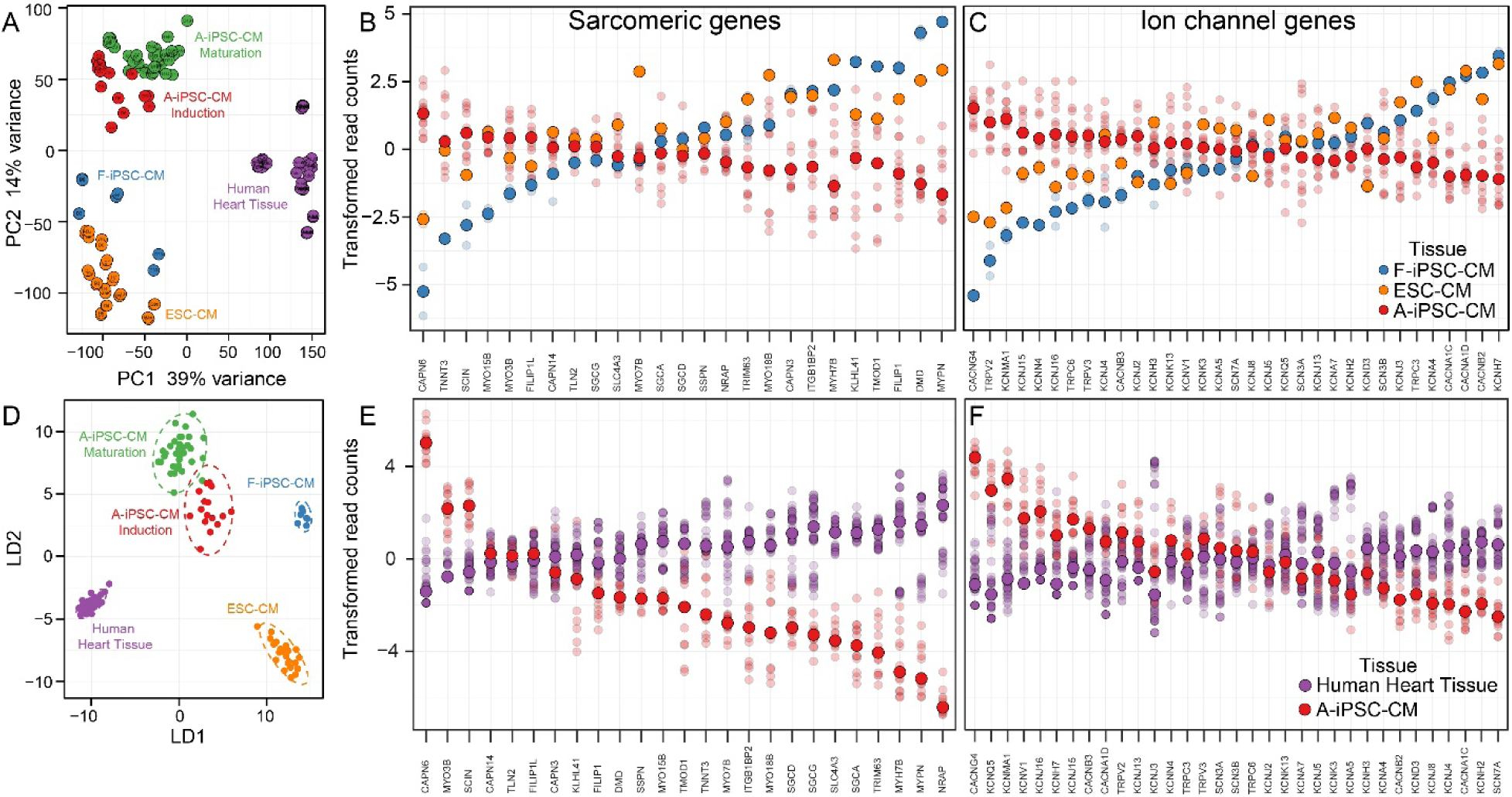
The distinct transcriptional signature of amniocytes-iPSC-CMs. (A): Principle component analysis of amniocyte-iPSC-cardiomyocyte induction (Day0, 3, 6, 9, N = 24), amniocyte-iPSC-cardiomyocyte maturation (Day0, 7, 14, 21, 35, 49, N = 36), fibroblast-iPSC-cardiomyocyte (F-IPSC-CMs Day0, 3, 7,30, N = 8), human ESC-Cardiomyocyte (ESC-CMs Day0, 1, 2, 3, 4, 5, 6, 8, 10, 15, 31, N = 26), and human heart tissue (purple, N = 48). (D): Linear discriminant analysis of amniocyte-iPSC-cardiomyocyte induction (Day0, 3, 6, 9), amniocyte-iPSC-cardiomyocyte maturation (Day0, 7, 14, 21, 35, 49), fibroblast-iPSC-cardiomyocyte (F-IPSC-CMs Day0, 3, 7, 30), ESC-CMs Day0, 1, 2, 3, 4, 5, 6, 8, 10, 15, 31), and human heart tissue (purple). RNA-seq scatterplot comparing upregulated (B) sarcomeric genes and (C) ion channels and in amniocyte-iPSC-CM (A-iPSC-CMs, Day 21), Fibroblast-iPSC-Cardiomyocyte (F-IPSC-CMs Day30, blue), human Embryonic Stem Cell-cardiomyocytes (ESC-CMs Day31, orange). RNA-seq scatterplot comparing upregulated (E) sarcomeric genes and (F) ion channels and in amniocyte-iPSC-CM (A-iPSC-CMs, Day 21, red) to human heart tissue (purple). Solid circles are the median value (bars ± standard deviation) for each group and the transparent circles are expression levels for individual samples.

Alternately, using the minicircle-LGNSO approach, while expression of the minicircle-encoded factors increased, the core pluripotent circuits did not significantly change, and no iPSC lines were generated. Minicircle-LGNSO contains a CMV promoter followed by the five genes LGNSO in the cassette. Based on the molecular action of this system, each of these genes should be expressed at equal levels, but we observed that Lin28, the first gene in the minicircle cassette, was expressed at extremely high levels, and subsequent genes show a step-wise lower level of expression (Sup Figure 7). In fact, Sox2 and Oct4 could not be detected above baseline levels in amniocytes. Low expression levels of these two key reprogramming factors may explain why amniocytes failed to convert into a pluripotent state. To examine whether forced expression of Lin28 and Nanog is sufficient for partial reprogramming, bypassing the baseline pluripotent state, and allowing these cells to differentiate directly into mesodermal precursors, amnioctyes were reprogrammed with the minicircle-LGNSO and then differentiated using cardiogenic protocols. Like previous manipulations, amniocytes did not increase expression of cardiac markers, and no spontaneous beating occurred. Taken together, these results indicate that amniocytes are resistant to reprogramming into a cardiogenic fate by a wide variety of methods (Sup Table 1 and 3), contrary to our original hypothesis.

**Figure 7.**
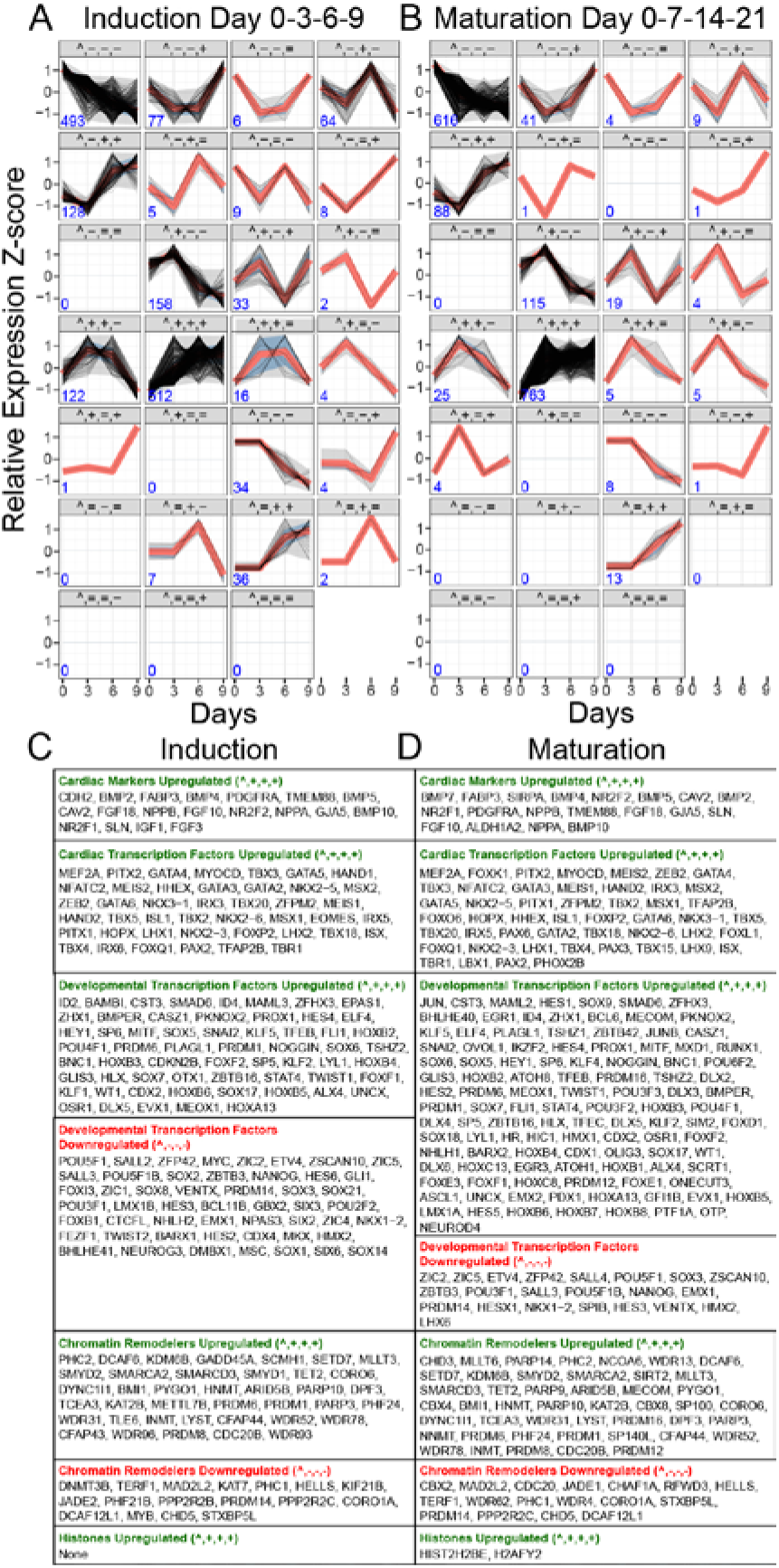
Identification of co-regulators during early induction and maturation. (A): Induction time course (Day 0, 3, 6, 9, N = 24) and (B) maturation time course (Day 0, 7, 14, 21, N = 36) of differentiating pluripotent iPSCs to beating cardiomyocytes. RNA-seq time course data clusters into 27 possible patterns, identifying temporally coordinated genes that are unchanged, activated or repressed. 1721 well-characterised genes, based on class, function, or pathway were processed by this clustering algorithm. The likelihood-ratio test was set at </= 1 and the Z-score threshold (relative expression) for change was 0.05. The red line indicates the median, the dark grey areas are 98% confidence interval and light grey are 95% confidence interval. The time point day 0 was sequentially compared to the induction day 3, 6, 9 and maturation day 7, 14, 21. The values were defined as (=) no statistical change, (-) decrease, and (+) increase between the two time points. N in lower left corner = number of genes that fell into each particular pattern. During (C) induction and (D) maturation time courses, groups of cardiac markers, cardiac transcription factors, developmental transcription factors, chromatin remodelers, and histones that shared a significant up or down changes in gene expression (padj < 0.01). Genes are listed in order from highest to lowest significant change.

### Amniocytes have non-conforming DNA methylation

DNA methylation may account for the strong suppression of important cardiogenic genes in amniocytes. DNA methytransferases and DNA demethylases are known to establish and maintain DNA methylation patterns. As a starting point, we examined the differences in expression of these factors in amniocytes, pluripotent cells, and heart tissue. Interestingly, amniocytes express a distinct pattern of DNA methyltransferases and DNA demethylases (Sup Figure 8). We next used genome-wide meDNA-seq and examined whether the methylation status could be manipulated by DNA hypomethylating agents, focusing particularly on cardiac-specific genes. Treatment of amniocytes with 5-Aza slightly reduced DNA CpG methylation levels on a genome-wide level (Sup Figure 9). However, for important cardiogenic genes, the DNA methylation levels did not change at promoter domains or across open reading frames (ORF). According to established models, increased levels of DNA methylation in promoter domains is linked to gene suppression. To examine whether this relationship holds true in amniocytes, we plotted the DNA methylation level of amniocyte transcriptional start sites (TTS, log10 mean of transcripts per million) versus gene expression levels for selected cardiogenic genes (1994 genes). These two variables do not fit in a linear fashion. Instead, the distribution is bimodal. A majority of TTSs were either hypomethylated or hypermethylated and genes were simultaneously expressed at high levels, densely populating the extreme ends of the plot (Sup Figure 10). To determine whether specific types of genes and their promoters express a similar bimodal pattern, 1994 cardiogenic genes were grouped into nine categories (Sup Table 3: Cardiac markers = 43, Cardiac transcription factors = 133, chromatin remodelers = 768, developmental transcription factors = 619, histones = 88, ion channels = 160, reference genes = 6, and sarcomeric factors = 174, Wnt signaling pathway = 130). Unexpectedly, all nine gene categories exhibited a bimodal pattern in methylation and expression at the promoter (Figure 2C). We also made histograms for the calculated average methylation values of selected gene TTS, sorted by binning TSSs expressed above 20 reads and below 20 reads (Sup Figure 11).

The bimodal distribution was still present for cardiogenic genes. To explore this relationship further, we made two separate sets of stacked barplots, and sorted by TSSs that were 1) hypermethylated or hypomethylated (Sup Figure 11) and 12) above 20 reads or below 20 reads (Sup Figure 13). For selected cardiogenic promoters or TTSs, DNA methylation and gene expression do not correlate in a linear fashion but exhibit a bimodal distribution.

Although the significance of the bimodal distribution is unclear, it suggests relatedness of functional groups or shared common pathways. In somatic cells, start sites are generally unmethylated, while gene bodies are sometimes methylated and this pattern depends on cell-type (Jones, 2012). To determine the if gene bodies also have a bimodal distribution, we assessed methylation status of selected cardiogenic genes in amniocytes. We applied a BSmooth algorithm to better visualise differentially methylated regions of cardiogenic genes (Hansen et al., 2012). After closely examining BSmooth plots for individual cardiac-relevant genes, we noticed that some genes have distinct methylation patterns (Sup Figure 14), particularly across the length of the open reading frame (Figure 2A and 2B). For example, some transcription factors like Gata6 are relatively hypermethylated across the gene body (Figure 2A), while in contrast, transcription factor Nkx2-5 is relatively hypomethylated across the gene body (Figure 2B).

To determine whether methylation of the gene body is a stronger predictor of gene expression than methylation of the promoter region, we generated a genome-wide scatterplot of gene expression for the full length ORF against DNA methylation (Figure 2D). Interestingly, the data is non-normally distributed, highly scattered and nonlinear (r value = -0.20). Of the four quadrants, hypermethlated gene bodies that are highly expressed contained a surprising number of data points (Quadrant I). The other three quadrants also contained a large number of data points: Quadrant II (hypomethylated and high expression), Quadrant III (hypomethylated and low expression), Quadrant IV (hypermethylated and low expression). Although the largest number of genes clustered into Quadrants I and IV, another smaller cluster was observed in Quadrant III, suggesting a bimodal relationship is present here as well.

To determine whether specific types of genes and their gene bodies have a predictable pattern in methylation and expression, we analyzed our nine gene categories (described above). The nine gene categories had a distinct pattern and exhibited a variable distribution of data points, differing among class, function, and pathway (Figure 2E). We calculated linear regression slopes, linear regression p-values and Pearson’s r for each of these nine scatterplots. To further explore whether other gene elements follow nonconforming patterns, we plotted methylation level between the TSS and ORF (Sup Figure 15). The pattern in this plot is striking, Quadrants I, II, and IV contained many data points, but Quadrant II contained very few. Based on these analyses, categories of genes appear to behave differently, suggesting a complex and varied relationship exists between DNA methylation and gene expression that is not easily explained, but may reflect specific gene-types. In particular, cardiac transcription factors, developmental transcription factors, and histones exhibit a non-random, bimodal pattern of clustering in specific areas of the plots (blue boxes with percentages, Figure 2E; see also Sup Table 4), suggesting an undescribed biological process is differentially demethylating exons.

Taken together, DNA methylation and gene expression in human amniocytes exists in a relationship that does not conform to standard models and that may contribute to a complex repressive mechanism that prevents cardiomyocyte differentiation directly from amniocytes.

### Amniocyte-iPSCs share a similar yet distinct baseline pluripotent state to cell-types derived from multiple sources

Since amniocytes are refractive to direct reprogramming, we investigated whether non-integrative virus-mediated pluripotent reprogramming was sufficient to overcome the strong transcriptional repression. Sendai viral OKSM reprogramming efficiently and quickly produced multiple Amniocyte-iPSCs lines. We confirmed pluipotency by examining pluripotent surface markers Tra-1-60 and SSEA4 (Figure 3A and 3B) as well as nuclear expression of core pluripotent regulators Nanog, Sox2, and Oct4 (Figure 3C-G) (Figure 3D-H). Sendai-generated amniocyte-iPSCs upregulated expression of core pluripotent regulators at much higher levels than the same untransduced patient control sample (Figure 3I). In the scatterplot of principle component analysis (Figure 3J), unmanipulated amniocytes express more PC2 variation than true pluripotent cells. A large pluripotent cluster containing many iPSC and ESC datasets gathered from several laboratories, including A-iPSCs, is distinct from unmanipulated amniocytes. Because pluripotent cells clustered together so tightly, gene variation in A-iPSCs appears to be very similar to variation in pluripotent cells derived in other laboratories. To better compare pluripotent cells, we excluded unmanipulated amniocytes, and reprocessed the PC analysis (Figure 3K). In general, the PC analysis of pluripotent cells derived from a variety of cell-types suggests A-iPSCs may have a baseline pluripotent state that is similar, but not identical to iPSCs and ESCs. Linear discriminant analysis (Figure 3L) of the top ten most variably expressed genes in A-iPSCs and pluripotent cells derived from other sources indicates A-iPSCs have a distinct pluripotent baseline state. To determine whether highly expressed embryonic stem cell repressors decrease as a result of reprogramming to a pluripotent state, we examined the expression level of 44 genes previously identified as potential transcriptional repressors that block direct transdifferentiation (Maguire et al., 2013). Many embryonic stem cell repressors markedly decreased expression in A-iPSCs (Figure 3M). Altogether, this data demonstrates that reprogrammed amniocyte-iPSCs appear to be cleared of strong transcriptional gene repression, likely by following standard pluripotent pathways.

### Amniocyte-iPSC-cardiomyocytes

Recent advances in iPSC-cardiomyocyte differentiation protocols allow efficient conversion to spontaneously contracting sheets of cardiomyocytes (CMs) (Burridge et al., 2014). We tested whether Sendai generated amniocyte iPSCs can be differentiated into cardiomyocytes using the CDM3 method, and found it produced vigorously contracting sheets (>120 contractions per minute) in 7-8 days (Sup Video 1). To examine cardiovascular differentiation, we analyzed expression of common cardiac markers at 7-10 days post differentiation. Spontaneously contracting amniocyte-derived iPSC-CMs (A-iPSC-CMs) co-express key cardiac regulators, sacromeric proteins and ion channel proteins of the human heart (Figure 4A-E). A-iPSC-CMs exhibit nuclear expression of Mef2c, Gata4, and Tbx5 as well as the cytoplasmic sarcomeric contractile proteins alpha-actinin, cardiac Troponin T (CTNT) and myosin heavy chain II (MF20) (Figure 4A-D). CTNT-positive cells co-express the cardiac ion channel Kcna5 (Figure 4E). Some amniocyte-iPSCs give rise to myofibroblasts or smooth muscle cells and stain SM22-positive, while others differentiate into epithelial cells and stain ZO1-positive (Figure 4 G, H, and J). Human ventricular cardiomyocytes stain positive for the same markers (alpha-actinin), but have different sarcomeric structure, organization, and morphology than immature A-iPSC-CMs (Figure 4J).

To determine whether A-iPSC-CMs acquire a transcriptional signature of human hearts, we performed RNA-seq analysis and compared the results to publicly available datasets derived from human hearts. Interestingly, differentiated A-iPSC-CMs express early-, mid-, and late-cardiac markers at similar levels and patterns as human hearts (Figure 4K). Expression levels for early markers are generally higher in A-iPSC-CMs, and mid- and late markers are generally lower in A-iPS-CMs. This expression profile analysis suggests that the cardiac signature of A-iPS-CMs is closer to a fetal phenotype than an adult phenotype. However, these results do not determine the atrial or ventricular characteristics of A-iPS-CMs.

To assess the subtype identity of A-iPS-CMs, we performed electrophysiological recordings in single cells at 20 days post onset of beating (Figure 5A and 5B), using the near-infrared fluorescent voltage-sensitive dye di-4-ANBDQBS to measure optical action potentials (Lopez-Izquierdo et al., 2014). To determine the ventricular-or atrial-like phenotypes using optical APs recorded using fluorescent dye, APD50 were calculated as the time at which repolarization of the optical AP achieved 50% of the optical amplitude (Figure 5A). The distribution of APD50 values and optical AP morphologies allowed us to distinguish two main subpopulations of CMs. Most notable, a majority of individual A-iPSC-CMs exhibited a ventricular-like action potential, defined by a clear plateau phase with longer action potential durations. Less frequently, atrial-like cells were identified, defined by a more triangular shape and the absence of a plateau phase. Intermediate cells were defined as sharing features of both triangular and plateau action potential morphologies. These analyses indicate that A-iPSCs efficiently differentiate into beating cardiomyocytes that are predominately ventricular-like in their electrical phenotypes.

To confirm the presence of a ventricular cardiomyocyte subtype, we stained contracting sheets for atrial and ventricular markers (Sup Figure 16). At Day 7 post differentiation, immunostaining images and qPCR analyses indicate the atrial marker MLC2A (Myl7) is strongly expressed, but not the ventricular marker MLC2V (Myl2). Although MLC2A (Myl7) is commonly used to label atrial tissue, it marks myocardial cells in both the atria and ventricles during development (Cai et al., 2003) and is markedly upregulated as part of the re-expressing the fetal program in human failing hearts (Depre et al., 1998; Dirkx et al., 2013), as confirmed in Sup Figure 17. Although we observed MLC2A (Myl7) expressed in A-iPS-CMs, this marker completely overlaps with the general cardiomyocyte marker CTNT and does not appear to be specific for atrial cardiomyocytes (Sup Figure 16). At later time points (Day20), the ventricle-specific marker MLC2V (Myl2) starts being detected by immunofluorescence (Figure 5F-G).

Interestingly, qPCR analysis indicates MLC2V (Myl2) expression peaks at Day 20, at subsequent time points Day 30 and Day 60, expression for this ventricular marker decrease (Figure 5I). Furthermore, expression of MLC2V (Myl2) seems to be dependent on protocol used to generate A-iPSC-CMs, different differentiation protocols generate A-iPSC-CMs expressing a broad range of MLC2V (Myl2) from low to high (Figure 5J). Taken together, reprogrammed amniocyte-iPSCs differentiate into *bona fide* cardiomyocytes and other cardiovascular cell-types with a predominantly ventricular physiologic profile. Based on phenotypic analyses of gene expression and electrophysiology, these studies suggest A-iPSC-CMs have a fetal-like cardiac nature, but also have a propensity to differentiate towards a ventricular-like cell-type. This might make them ideal candidates for cell therapies of ventricular CHDs.

### Amniocyte-iPSC-cardiomyocytes exhibit a distinct transcriptional signature

Independent studies have generated fully functional cardiomyocytes from a variety of different somatic cell sources and pluripotent cell lines (Burridge et al., 2014; Lan et al., 2013; Lian et al., 2012; Tohyama et al., 2013; Zhang et al., 2009; Zwi et al., 2009). It is not known whether cell-type of origin or differentiation protocol has a greater influence on final cardiomyoctye phenotype. It has also been proposed, as long as the cardiomyocytes generated are functionally normal, slight differences in gene expression are less important. However, gene expression can have profound impact on both form and function. To address similarities and differences in cardiomyocytes generated from different studies in terms of overall internal structure of genome-wide datasets, we compared amniocyte-derived cardiomyocytes to fibroblast- and ESC-derived cardiomocytes (Sup Figure 18). Importantly, both the state of pluripotency and the differentiation approach/protocol appear to influence cardiomyocyte differentiation (Figure 6A and D). Principle component analysis and linear discriminant analysis reveal the transcriptomes of different cardiomyocyte differentiation time courses are quite distinct. Only the current induction and maturation datasets had pluripotent cells overlap in the starting position at Day0.

This should be expected because these experiments were performed in the same laboratory, albeit at different times. Although all three of these protocols produce contractile cardiomyocytes, the different datasets maintained separation or even slight divergence as they differentiate along cardiovascular pathways. It is important to note that A-iPSC-CMs were unpurified, while Fibroblast-iPSC-Cardiomyocytes (F-IPSC-CMs) were purified in glucose-depleted, lactate supplemented conditions (Ang et al., 2016), and human Embryonic Stem Cells-Cardiomyocytes (ESC-CMs) were purified using Percoll density gradients. The complete results of all PC 1-10 combinations (45 total) are presented in the supplementary information (Sup Figure 19). It is also important to note, in plot PC1 versus PC2, A-iPSC-CMs and ESC-CMs datasets are most dissimilar, but in a number of other PC combinations, the A-iPSC-CMs and ESC-CMs datasets are nearly superimposed, and positioned far away from the F-IPSC-CMs dataset. This observation suggests there are groups of genes driving these similarities and differences, and different PC plots recognise these patterns. Confounding technical factors related to generating and processing RNA-seq data should be considered. We performed a more in-depth comparison of the A-iPSC-CMs and F-IPSC-CMs datasets (Sup Figures 20-24). These results suggest the starting point of pluripotent cells and protocol/pipeline-specific differences can greatly bias the ending point of terminally differentiated cardiac cells.

To better define the status of terminally differentiated A-iPSC-CMs, we examined a select group of ion channel, sacromeric- and myofibril-related genes as a reflection of function. RNA-seq time course analysis identified 34 ion channels and 53 sarcomeric proteins that are specifically upregulated in pluripotent-to-cardiomyocyte conversion (Figure 5B, 5C, 5E, 5F).

Although the general pattern of A-iPSC-CMs overlap with other pluripotent-to-cardiomyocyte cell types, there are significant differences, implicating potential differences in electrophysiologic and contractile function. As a molecular measure of cardiomyocyte function, these results demonstrate that expression of developmentally and cell-type specific ion channels, sacromeric- and myofibril-related genes are distinct among cardiomyocyte derived from different cell populations.

### Cardiogenic networks of coordinated gene expression

A complex transcriptional program coordinates cardiovascular developmental decisions (Wamstad et al., 2012). Based on mathematical models, different types of cluster algorithms have provided insight into gene functions and networks because genes in similar groups tend to be functionally related (Kim et al., 2015; Si et al., 2014). To visualise common patterns of global gene expression in our pluripotent-to-cardiomyocyte differentiation time courses, we employed a self-organizing map (SOM) approach to analyze both induction and maturation datasets.

Although we performed this analysis genome-wide, to make the datasets in a more manageable size, we narrowed the genes down to 1721 factors known or implicated to play a role in cardiogenesis. For simplicity, we determined the total number of possible patterns (27) across four time points (Sup Figure 25 and Figure 6A and 6B). The cluster pattern approach properly placed well-characterised cardiogenic factors in expected patterns of gene expression. In tandem, we performed differential gene expression analyses at each time point to confirm which factors were significantly upregulated or downregulated (Sup Figures 26-31). For example, the key core cardiogenic transcriptional factors Mef2c, Tbx5, Nkx2-5 and Gata4 as well as sarcomeric proteins Ctnt, Myh6, Myh7 and alpha-actinin also fell into expected patterns and are upregulated. In our selected group of 1721 genes, many developmental transcription factors not known to play a role in cardiac development were upregulated. The transcription factor families include basic helix-loop-helix, basic leucine zipper, forkhead boxes, aATA zinc finger domain containing, helix-turn-helix ETS type domain containing, HOXL subclass homeoboxes, NKL subclass homeoboxes, POU class homeoboxes, PRD class homeoboxes, SINE class homeoboxes, SMAD family, SRY (sex determining region Y)-boxes, ZF class homeoboxes, and Zinc fingers, C2H2-type. Conversely, pluripotent genes are markedly downregulated and also cluster in expected patterns (Figure 7C, developmental transcription factors downregulated).

Importantly, our clustering algorithm identified numerous putative co-regulators of cardiomyocyte differentiation (Figure 7C Induction and 7D Maturation, and Sup Table5 full list of genes). These results provide a detailed catalogue of novel clustering relationships and potential co-regulators that may play important roles in cardiomyocyte differentiation. Although the number of possible cluster patterns is the same for the initiation and maturation datasets, the genes that fell into each pattern differed, suggesting some co-regulators play dual roles or carry heavier weight during important steps of functional maturation.

## DISCUSSION

Using current induction methods, second trimester amniocytes rarely differentiate into functional cardiomyocytes. This study offers insight into repressive mechanisms at play in restricting the cardiogenic potential of amniocytes. Despite evidence of being poised, our data indicates the amniocyte DNA methylome interconnects with the transcriptome in nonconforming ways to repress known cardiac inducers. This distinct molecular phenotype makes expandable amniocyte cell lines strongly resistant to directed cardiomyocyte differentiation, but iPSC reprogramming easily removes this transcriptionally repressed state. The time-course analysis of reprogrammed and differentiated A-iPSC-CMs identified temporal co-regulators of induction and maturation belonging to diverse gene families of cardiac transcription, developmental transcription, and chromatin remodeling. Additionally, comparative transcriptomics analyses reveal A-iPSC-CMs exhibit a distinct sarcomeric phenotype and expression of some ion channel genes not previously reported to play a role in development of the cardiac action potential. We conclude that overcoming the powerful repressive mechanisms of DNA methylation and transcription are critical when converting amniocytes into fetal cardiomyocytes.

Interestingly, global methylation levels in amniocytes are quite distinct from other extraembryonic cell-types [66]. In our study, cardiomyocyte-specific genes and other classes of genes exhibit distinct patterns of DNA methylation, but the functional significance of these patterns is not known. Through the examination of 125 human cell lines, the ENCODE Project presents strong evidence for the general rule: increasing methylation levels decreases chromatin accessibility [67]. Comprehensive maps and large datasets provide valuable information about general rules. However, distinctions might be overlooked when analyzing large genome-wide datasets. While the amniocyte DNA methylome follows some general rules, this cell-type has many surprising non-conforming features, which appear be heavily influenced by particular pathways.

The dynamics between DNA methylation and gene expression have not been defined in amniocytes. Developmental stage and tissue-type are known to influence the cardiac methylome (Gilsbach et al., 2014; Liang et al., 2011; Tompkins et al., 2016). DNA methylation analyses from these studies support the widely held model that the DNA methylation status of these regions strongly correlate with gene expression. Cardiomyocyte transcriptional start sites have an inverse relationship, while gene bodies have a positive relationship. In contrast, our results indicate that amniocyte DNA methylation and gene expression only weakly correlate, and when common functional groups or pathways are analyzed separately, the relationships can be positive, negative or uncorrelated, implying diverse mechanisms regulate different types of genes. In line with our bioinformatics analyses, one previous study documented markedly different DNA methylation patterns for cardiac structural genes compared with cardiac-specific transcription factors (Gu et al., 2014). We propose that these two cardiomyocyte-specific gene categories (e.g. structural versus transcription) may be part of separate functional classes controlled by distinct, but unknown, pathways. We propose that specific methylation patterns on amniocyte DNA, specifically on key transcription factors, may play an important role in suppressing cardiomyocyte transdifferentiation.

While the exact origin of human amniocytes is not known, morphological and molecular analyses in animal and human amniotic fluid cells suggest amniocytes originate from multiple fetal epithelial tissues (Dobreva et al., 2010). It is therefore surprising amniocytes seem to possess a poised cardiogenic potential. It has been proposed amniocytes have a higher degree of transcriptional plasticity than other lineage restricted cell-types. Contrary to our initial hypothesis, we find that amniocytes are resistant to direct differentiation into cardiomyocytes using established protocols. Our findings suggest amniocyte transcriptional plasticity into the cardiomyocyte cell fate is restricted.

Like most terminally differentiated somatic cell-types, induced pluripotent stem cell reprogramming removes transcriptional repression in amniocytes. Reprogramming experiments demonstrate that Sendai-OKSM returns amniocytes to baseline pluripotency, but their status is distinct from other iPSC lines derived across disparate cell-types and tissues. The distinct pluripotent signature of amniocytes may contribute to a biased differentiation potential. The immunostaining and electrophysiologic results indicate that amniocyte-iPSC-cardiomyocytes bias toward a ventricular phenotype, which could be useful for some tissue replacement approaches.

Although we used a different RNAseq analysis pipeline and a different starting cell source, our general findings in transcriptome dynamics are consistent with previous studies in mouse (Wamstad et al., 2012) and human cardiomyocytes (Paige et al., 2012; Tompkins et al., 2016). Previous studies propose pluripotent-derived cardiomyocytes are phenotypically and transcriptionally related to fetal hearts (Burridge et al., 2014; van den Berg et al., 2015). Despite the similarities, in this study, cardiac marker gene expression in fetal hearts is quantitatively closer to children and adult hearts than to amniocyte-iPSC-cardiomyocytes. This observation suggests important signaling factors of normal heart development are missing with *in vitro* culture systems.

A genetic map of inductive and maturational signaling factors controlling cardiomyocyte and nonmyocyte populations of the heart is incomplete. Longitudinal RNAseq analyses in pluripotent-derived cardiomyocytes are helping close the gap. Our principal component and differential gene expression analyses demonstrate pluripotent-derived cardiomyocytes do not match the transcriptome of human heart tissue, but seem to be within striking distance. The initiation (Days 0, 3, 6, 9) and maturation (Days 0, 7, 14, 21, 35, and 49) time courses exhibited overlapping transcriptome trajectories. However, in our maturation time course, once A-iPSC-CMs reached day 21 post-differentiation, they developmentally stopped maturing. This is consistent with results in mouse ESC-derived cardiomyocytes (Li et al., 2015). It is unknown what signaling factors and conditions are required for maturation to proceed. The next phase in culture-derived cardiomyocytes reprogramming technology will be to understand the signals and mechanisms promoting the maturation of cardiomyocytes to share similar chamber-specific and cell-type-specific phenotypes with fetal, child, adolescent, and adult hearts.

## MATERIALS AND METHODS

### Human amniocytes

ARUP laboratories (Salt Lake City, Utah) provided whole amniotic fluid samples, 15-30 gestational weeks. 494 patients contributed 100µl of amniotic fluid to this study. When cultured, approximately 25% of these samples derived adherent, expandable amniocytes. University of Utah IRB determined oversight was not required (IRB_00040970) because de-identified amniocytes do not meet the definitions of Federal regulations on Human Subjects Research. Amniocyte isolates that expanded in culture were selected for iPSC reprograming if they were genetically normal. ~100ul of whole amniotic fluid was added to 6ml of AmnioMAX-II Complete Medium (ThermoFisherScientific: 11269016) supplemented with 10% Embryonic Stem Cell-qualified Fetal Bovine Serum media (ThermoFisherScientific: 10439016) or 10% EquaFETAL FBS (Atlas Biologicals: EF-0500-A). After seven days, media was changed. Once isolates were identified as growing, cells were passaged 1:1, 1:3, and 1:6. Fully confluent 6-well plates were cryopreserved in Recovery Cell Culture Freezing Medium (ThermoFisherScientific: 1264810). Primary amniocyte cell lines generated for this study were not authenticated. Since these lines were derived from clinical samples, a small number of samples would arrive contaminated with fungus. Fungal contaminated samples were excluded from the study.

### Human ventricular heart tissue

Serving as controls for cardiomyocyte morphology and gene expression, de-identified heart samples were obtained from the laboratory of Dr. Stavros Drakos. Small frozen pieces of tissue were thawed and prepared for immunostaining and imaging or homogenized for RNA isolation.

### Derivation of Sendai viral induced pluripotency in amniocytes

Early passage (1-5) human amniocytes isolated from amniotic fluid were expanded to plastic 6-well plates in AmnioMAX-II Complete Medium. To determine whether amniocytes could be tranduced by Sendai virus, we used the CytoTune-EmGFP Sendai Fluorescence Reporter (ThermoFisherScientific: 12558011: A16519). For reprogramming amniocytes, 6-well plates were grown to 50-80% confluency and then were transfected with CytoTune-iPS 2.0 Sendai Reprogramming Kit (ThermoFisherScientific: A16517), using the Feeder-Independent Protocol. To confirm the presence of iPSC colonies, we followed the live staining protocol and used mouse anti-Tra1-60 antibody (ThermoFisherScientific: 41-1000). When iPSC colonies grew to sufficient size and had a flat circular morphology, they were physically picked with a P200 pipette and moved to new wells for expansion. Alternatively, to improve recovery of colonies, we passaged the entire well with a modified Accutase Protocol. E8 media was aspirated, and 1ml of warm Accutase Solution (MilliporeSigma: A6964-100ML) was incubated at 37C for minutes.

Cells were scraped off with a cell scraper and 2ml of warm KO DMEM (ThermoFisherScientific: 10829018) was added to each well and media/cell were moved to a 15ml conical tube. Cells were spun down in15ml tubes at 160g (1300rpm) for 3 minutes. Media was aspirated and 1ml of E8 media treated with ROCK Inhibitor Y-27632, 10µM (Selleckchem: S1049) was added to the tube and cells were plated on vitronectin-coated 6-well plates (ThermoFisherScientific: A14700). A-iPSC colonies were passaged 5-10 times before they were cryopreserved. To induce early passage amniocytes into pluripotency, we followed the methods outlined in Invitrogen’s Cytotune-iPS 2.0 Sendai Reprogramming Kit (ThermoFisherScientific: A16517). In pilot studies, amniocytes were also reprogrammed using lentiviruses expressing OKSM (Riedel et al., 2014).

### Differentiation of A-iPSC-CMs

To differentiate A-iPSCs into cardiomyocytes, we modified the CDM3 protocol developed by Burridge et al (Burridge et al., 2014). Briefly, A-iPSCs were expanded in 6-well plates containing Essential 8 medium (ThermoFisher Scientific: A1517001) and grown to 75% confluence. On Day0, 100% E8 media was switched to E8/CDM3 (17%/83%) and treated with CHIR99021 3µM (MilliporeSigma: SML1046-5MG), a potent Wnt agonist. On Day1 (24hrs), E8/CDM3 (9%/91%) media was treated with CHIR. On Day2 (48hrs), E8/CDM3 (9%/91%) media was treated with 2µM C59 (Selleckchem: S7037), a Wnt inhibitor. On Day3 (72hrs), E8/CDM3 (9%/91%) media was treated with C59. On Day4 (96hrs), C59 inhibition was stopped and E8/CDM3 (9%/91%) media was changed. On Day5 (120hrs), E8/CDM3 (9%/91%) media was changed without manipulating Wnt-signaling pathways. On Day6-7 (144-168hrs), 100% CDM3 media was changed daily. Beyond Day7, CDM3 media was changed every other day.

### Immunohistochemistry, Confocal Microscopy, and Image Processing

Baseline amniocytes, A-iPSC, A-iPSC-CMs, and human heart samples were stained, imaged, and processed as previously described (Maguire et al., 2013). Staining for nuclear factors and cardiac structural markers was performed on permeabilised cells, while cells stained for pluripotent surface markers were unpermeabilised. For a complete list of primary antibodies used in this study: see Sup Methods Table 1. The following antibodies were purchased from the Developmental Hybridoma Studies Bank (DSHB) at the University of Iowa: 1) Solter, D. and Knowles, B.B deposited MC-813-70 (SSEA-4); 2-5) Jessell, T.M. and Brenner-Morton, S. deposited 39.3F7, 39.4D5, 40.2D6, 40.3A4 (Islet-1); 6) Schiaffino, S. deposited RV-C2 (troponin T, cardiac); 7) Fischman, D.A deposited MF20; 8) Greaser, M.L. deposited 9D10 (titin); and 9) Schiaffino, S. deposited TI-1 (troponin I, cardiac).

### qPCR

Total RNA was isolated using Ambion’s PureLink RNA Mini Kit (ThermoFisher Scientific: 12183018A) and measured on a NanoDrop 2000c spectrophotometer (ThermoFisher Scientific: ND-2000C). cDNAs were made using the SuperScript VILO cDNA Synthesis Kit and Master Mix (ThermoFisher Scientific: 11754050), 10 µM SYBR Select Master Mix (ThermoFisher Scientific: 4472908) and primer sets (designed by ThermoFisher’s OligoPerfect Designer, made by Operon's PCReady PCR & Sequencing Primers; see Supplemental Methods Table 2). PCR reactions were run with reference genes GAPD-Human (Roche: 05190541001), SDHA-Human, RPLO1-Human. On a LightCycler 480 instrument (Roche), the absolute quantification/2nd Derivative Max setting generated raw Cp values and was used to calculate gene expression levels.

### RNA-seq

RNA isolated as described above was used for both qPCR and RNA-seq analysis. Bioanalyzer 2100 (Agilent Technologies) was used to verify RNA integrity. 1.5 µg of total RNA from each sample was converted to mRNA-seq library at the using TruSeq Stranded mRNA Library Prep Kit (Illumina). Libraries were sequenced on a HiSeq 2500 (Illumina), containing multiple barcoded samples per lane. Sequences were aligned to human genome build hg19 using Star (Dobin et al., 2013). For each dataset, normalised count matrix underwent a variance-stabilizing transformation using the DESeq2 package (Anders and Huber, 2010) in R (Team, 2012). Expressed or not expressed cutoff was set at 20 reads. Adult fetal and adult heart RNAseq datasets were downloaded from Gene Expression Omnibus (GEO).

### Bisulfite sequencing

For 24 hours on 6-well plates, confluent amnioctyes were untreated (DMSO alone) or treated with different combinations of hypomethylation agents: 5-azacytidine 10μM, RG108 1μM, or AMI-5 5μM, dissolved in DMSO or molecular grade water. DNA was extracted from approximately 1-2 million cells per confluent sample, having a target concentration of 100ng of DNA and purified using a Qiagen non-organic DNA clean up kit. A focused-ultrasonicator (Covaris, AFA Technology) sheared the DNA. We spiked the sheared DNA with 1% unmethylated lambda DNA. 10 DNA samples underwent bisulfite treatment (Qiagen, Epitect bisulfite kit, Cat#59104) and sequencing on four lanes of HiSeq 2000 (Illumina), generating 101bp paired end reads. 17 Fastq files from four different patients were aligned to Novoalign in bisulfite mode (human genome build hg19) and BAM files were generated. Useq software (Nix et al., 2008) in R (Team, 2012) created count matrices that overlapped the annotated genes (Ensgene annotation) for 1) coverage and 2) HcG methylation. Replicates were merged and the coverage was either 4x low or 10x high. To visualise individual genes of interest in our WGBS dataset, the coverage was smoothed using a Bsseq R-package in Bioconductor (https://bioconductor.org/). The BSmooth program generated high-resolution plots of DNA methylation fractions (Hansen et al., 2012).

### Electrophysiology

Methods for optical AP recordings and fluorescent-based optical action potential imaging have been previously described (Lopez-Izquierdo et al., 2014). Briefly, A-iPS-CMs at 20 days post onset of beating were plated at low density to perform single cell recordings, coverslip plated witn A-iPS-CMs were loaded with di-4-ANBDQBS in buffer physiological solution containing (in mmol/l) 126 NaCl, 4.4 KCl, 1.1 CaCl2, 1 MgCl2, 11 glucose, and 24 HEPES (pH 7.4 with NaOH); which was prepared daily from a stock solution of di-4-ANBDQBS dissolved in ethanol. Incubated cells in 20 µM di-4-ANBDQBS for 2-5 min resulted in fluorescent signals with excellent signal-to-noise ratios and minimal cellular toxicity for the duration of the recordings.

After incubation, coverslip plated witn A-iPS-CMs were placed on the stage of an inverted microscope, where the fluorescent voltage-sensitive dye was washed out by perfusion with control buffered solution. A-iPS-CMs loaded with di-4-ANBDQBS were excited by a 200-m W red solid-state laser (660 nm) coupled, fluorescence signal was recorded by an electron multiplied (EM) charge-couple device (CCD) camera (iXon 860, Andor Technology, Belfast, UK) connected to the video port and equipped with a 700 nm long-pass filter (Omega Optical). Cells were paced at a fixed CL of 800 ms to eliminate any variation in baseline. All experiment were performed at 36-37°C.

## DATA AVAILABILITY

Access to MeDNAseq and RNAseq data sets are listed in the supplemental information.

### ACKNOWLEDGEMENTS

We thank ARUP laboratories (Department of Cytogenetics and Genomic Microarray and Institute for Clinical and Experimental Pathology) for providing human amniocyte samples for this study. We thank Benoit Bruneau, Jianhua Zhan and Timothy Kamp for providing cardiomyocyte differentiation protocols before publication, and Michael Riedel, Scott Cho, and Chuanchau J. Jou for teaching us the basics in reprogramming human induced pluripotent stem cells. We thank Nikos Diakos and Stavros Drakos for providing us with de-identified human ventricular tissue as controls for immunostaining and qPCR. We thank Yen-Sin Ang and Deepak Srivastava for providing their F-IPSC-CMs RNA-seq datasets before publication for comparison to our datasets. We thank Sharan Paul and Warunee Dansithong for helpful discussions on reprogramming. We thank members of the Yost Research Group for helpful discussions of results presented here, especially Jonathon Hill, Brent Bisgrove, Todd Townsend, Luca Brunelli.

